# Long-term Learning Induces Plastic Changes in Frontostriatal Circuits

**DOI:** 10.64898/2026.06.24.734256

**Authors:** Diancun Xuan, Diana C. Burk, Ramon Bartolo, Xiaoli Li, Bruno B. Averbeck, Hua Tang

**Author notes:** Correspondence (H.T.).

## Abstract

Neural activity in frontal-striatal circuits underlies reinforcement learning. Traditional theories suggest that reinforcement signals, which drive learning, strengthen connections within the basal ganglia. This strengthening is believed to shift information processing from cortical regions to subcortical regions as learning becomes established over time. To examine this hypothesis, we trained macaques to associate multiple sets of images with their values. Selecting different images led to either an increase (+2, +1) or a decrease (−1, −2) in the number of tokens, which subsequently determined the amount of juice reward the macaques received. We simultaneously recorded neuronal activity from orbitofrontal cortex, ventral striatum, amygdala, and dorsomedial thalamic nucleus, analyzing the dynamic changes in these brain regions during both the initial learning and overlearned stages. The results indicated that as learning progressed from the initial stage to the overlearned stage, information processing shifted from the ventral striatum to the orbitofrontal cortex, corresponding to the abstraction from stimulus value to state value. This finding challenges traditional theories and provides a new perspective on the neural circuit mechanisms of learning.

## Introduction

Reinforcement learning (RL), or the ability to adapt behavior in response to environmental feedback, is fundamental to cognitive flexibility. Extensive work has shown that RL is governed by frontostriatal circuits ^1–4^. For the learning of action-outcome associations, prior research has focused on the transition from goal-directed actions that are sensitive to outcome degradation to automated stimulus-response habits that are insensitive to outcome degradation ^5,6^. However, the neural dynamics of these processes during the transition from initial acquisition to overlearned associations remain less thoroughly explored.

During early learning, values associated with choices are updated, possibly through plasticity in frontal-striatal and amygdala-striatal circuits ^7–11^. Although the ventral striatum (VS) has traditionally been regarded as the primary site for reinforcement signals and rapid learning ^12,13^, recent evidence suggests that prefrontal signals, particularly from the orbitofrontal cortex (OFC), may precede or regulate striatal activity to bias motivational learning ^4,14^. During these early stages of learning, the OFC and amygdala (AMY) jointly evaluate contingent outcomes and guide initial choice selection ^15^.

Studies have suggested that as learning progresses into the overlearned stage, the goal-directed system goes through functional reorganization. Traditional “cortical-to-subcortical” shift hypotheses have proposed that subcortical structures, such as the basal ganglia, take over control as tasks become well-learned ^7,16–21^. However, recent research shows that the prefrontal cortex does not relinquish control but instead develops more stable, high-dimensional, and resilient representations ^22–28^. Consequently, long-term learning appears to transform prefrontal cortex representations, suggesting that expertise within a goal-directed framework is associated with more sophisticated cortical coding and consolidation ^29–31^. Despite these advances, the precise evolution of information processing across frontostriatal nodes during this transition remains insufficiently understood.

In this study, we investigated the functional reorganization of the frontostriatal circuit across different learning stages. Macaque monkeys were trained on a two-armed bandit learning task using a token-based symbolic reward system, with both novel and familiar blocks to assess learning and overlearning within a single session. Multi-region neuronal recordings were conducted simultaneously in the OFC, VS, AMY, and mediodorsal thalamus (MDt) to track the evolution of information processing. Contrary to the prevailing view that cortical involvement decreases as behavior becomes overlearned, our results reveal a shift in information processing from the VS to the OFC during the transition from early to late learning. Specifically, while the VS leads initial value updating, the OFC ultimately dominates stable state value coding and transmits stronger value signals to the VS during overlearning. Our findings provide a new perspective on how the frontostriatal circuit maintains and optimizes goal-directed representations over long timescales.

## Results

Two rhesus monkeys were trained on a two-armed bandit task. In this task, the monkeys collected tokens, which were periodically exchanged for juice rewards (Fig. 1a-b). In each block, we introduced 4 images corresponding to different token outcomes: +2, +1, −1, and −2. In each trial, 2 of the 4 images were presented on the screen. We defined a trial’s condition as a combination of image values, resulting in a total of 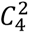 = 6 distinct conditions. The choice of an image led to the corresponding change of token with a probability of 75%, and no token change with a probability of 25% (Fig. 1b). Tokens were accumulated across trials and were cashed out every 4 to 6 trials, with one drop of apple juice for each token.

**Fig. 1.**
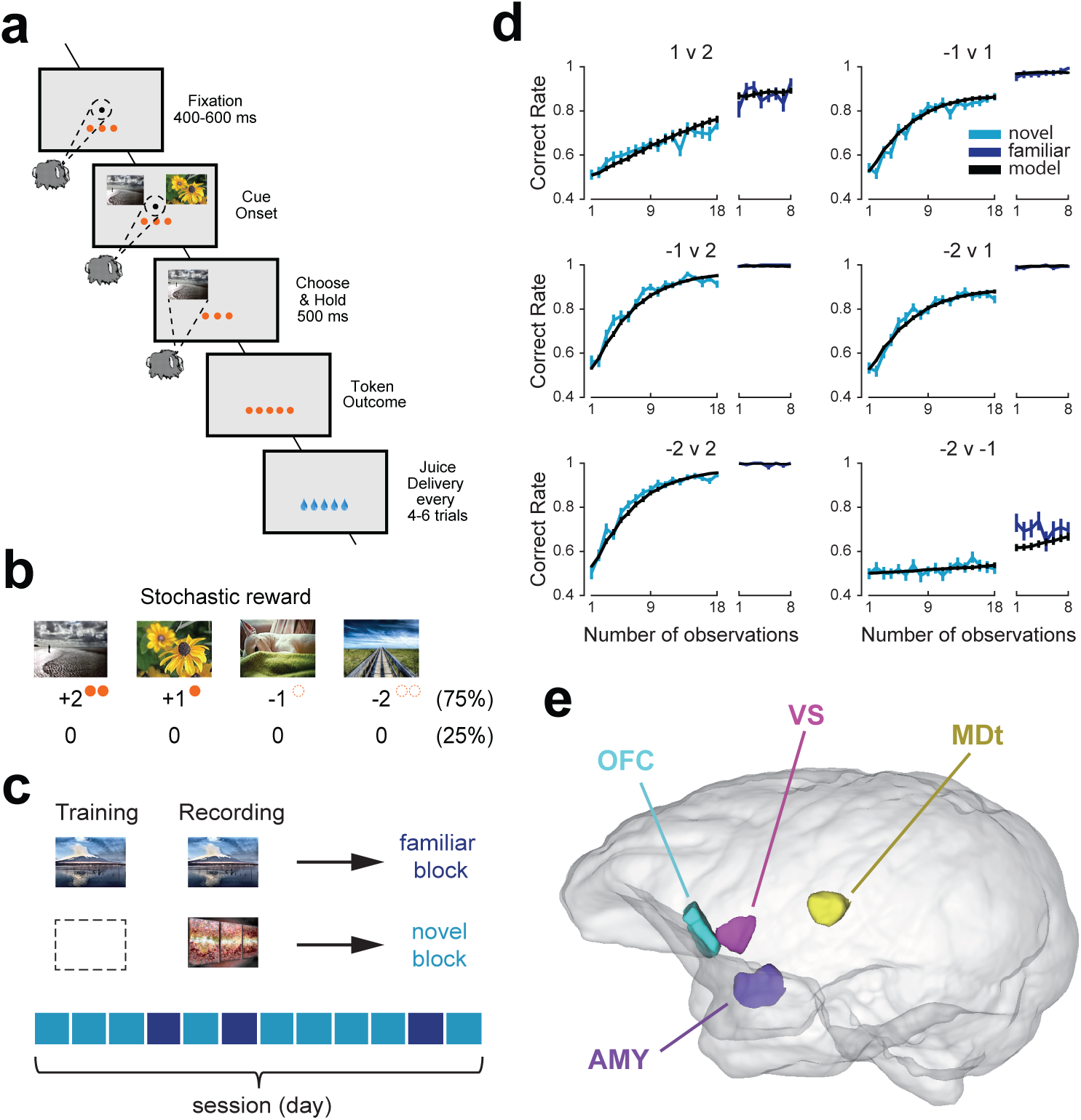
Task, behavior, and recorded regions. **a-c,** Behavioral task. **a**, Structure of an individual trial. Successive frames illustrate the sequence of events. In each trial, only 2 of the 4 images were presented. Monkeys chose between them and gained or lost (stochastic: 75% change, 25% no change) the corresponding number of tokens. Choices could be made as soon as the images were shown. Accumulated tokens were cashed out every 4 to 6 trials, with 1 drop of juice per token. **b**, In each block, the monkeys learned the values (+2, +1, −1, −2) of 4 novel images (an image set). **c,** Novel blocks and familiar blocks. Image sets in familiar blocks were learned during the training period, whereas those in novel blocks were never seen by the monkeys. A session (a day) comprises 9 novel blocks and 3 familiar blocks, with their order randomized. **d,** Choice behavior. Correct rate corresponds to the fraction of choosing the image with a higher value in each condition. Cyan and blue lines represent monkey behavior in novel and familiar blocks, respectively, and black lines represent the predictions of the RW model. Error bars represent mean ± SEM. Results were averaged across two monkeys (n = 50 sessions). **e,** Anatomical locations of simultaneously recorded brain regions: OFC (cyan), VS (magenta), AMY (purple), and MDt (gold).

There were 2 kinds of blocks, novel and familiar (Fig. 1c), based on the monkeys’ experience with the images. In novel blocks, novel images were presented to the monkey, and it had to learn the values of the images by choosing them and observing the outcome. Each novel block comprised 108 trials, in which each condition appeared 18 times (18 * 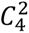 = 108). During task training, before neurophysiology recordings, some image sets were presented repeatedly every day for about 1 year. This allowed monkeys to overlearn the stimulus-outcome associations. During recording, these familiar image sets were presented in familiar blocks, and the monkey only needed to recall the values of each image to choose the better one. Each familiar block comprised 48 trials, in which each condition appeared 8 times (8 * 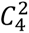 = 48). Discrimination of novel and familiar blocks provides a basis for comparing the learning of new stimulus-outcome associations with that of overlearned stimulus-outcome associations. Because blocks of each condition are presented interleaved (Fig. 1c), we can compare activity within the same session in the two conditions.

### Variability of choice behavior across learning stages

We quantified performance as the fraction of times the monkeys chose the image associated with the higher value in each condition (e.g., choosing +1 when presented with −1 and +1 images), hereafter called correct rate (CR). To describe the learning process, we regard CR as a function of the number of observations per condition (Fig. 1d).

We fit a multi-way ANOVA model to quantify the effects of different task variables on CR. In novel blocks, CR was significantly modulated by the number of observations (*F*_17, 45111_ = 111.25, *p* < 0.001), condition (*F*_5, 45111_ = 641.74, *p* < 0.001), and the number of tokens before choice (*F*_10, 45111_ = 12.36, *p* < 0.001). In familiar blocks, CR was not affected by the number of observations (*F*_7, 7033_ = 0.65, *p* = 0.714), but condition (*F*_5, 7033_ = 267.19, *p* < 0.001) and the number of tokens before choice (*F*_10, 7033_ = 2.42, *p* = 0.007) had significant effects on CR. Condition indicates the values of the presented images that lead to token updates, and the number of tokens indicates the accumulated value. The results indicate that, in novel blocks, the monkeys understood the meaning of tokens and adjusted their behavior to obtain more tokens as learning progressed. In familiar blocks, the monkeys’ performance had reached a stable limit, and no more progress was made. We also fit a Rescorla-Wagner (RW) reinforcement learning model to the choice behavior (Fig. 1d), as previously explored ^32^. It is used below to construct a partially observable Markov decision process (POMDP) model.

To compare the performance in different learning stages, especially between novel and familiar blocks, we conducted a proportion test. We split the trials into 3 equally sized stages: Early for 1^st^–48^th^ trials in novel blocks, Late for 61^st^ −108^th^ trials in novel blocks, and Overlearned for all the 48 trials in familiar blocks. Each pair of the 3 stages showed a significant difference (chi-squared proportion test, all *p* < 0.001), indicating further learning in familiar blocks than in late novel blocks.

### Neurophysiological recordings

Neural activity was recorded simultaneously from 4 regions within the ventral cortico-striatal-thalamo-cortical circuit using multi-site linear probes. We isolated 606 neurons from the orbitofrontal cortex (OFC), 829 from the ventral striatum (VS), 1,607 from the basolateral amygdala (AMY), and 1,035 from the medial portion of the mediodorsal thalamus (MDt). The semi-chronic recording procedure was described in detail in our previous work ^33^. The novel block data were previously analyzed to examine the neural coding of the token reinforcers ^33^. Here, with additional familiar block data, we aim to examine the plasticity of the neural circuit during learning.

### Neural activity changes with the learning process

We first examined how the strength of single neuron activity within a trial changed across the learning process. Different regions showed distinct activity gradients, most notably around the choice hold period and after the token outcome (Fig. 2a). To test this quantitatively, we split the trials into 3 stages: Early, Late, and Overlearned (as described earlier), and extracted activity around two time periods: Cue and Outcome, each corresponding to a 500 ms window (see Methods). We performed separate ANOVAs for Cue/Outcome periods and the 4 regions, focusing on the effect of learning stage (Fig. 2b). Specifically, the main effect of learning stage in the OFC was significant in the Cue period, but absent in the Outcome period (Cue, *F*_2, 1815_ = 5.93, *p* = 0.003; Outcome, *F*_2, 1815_ = 2.47, *p* = 0.085). In contrast, VS exhibited a significant effect of learning stage only during the Outcome period (Cue, *F*_2, 2463_ = 0.01, *p* = 0.993; Outcome, *F*_2, 2463_ = 4.51, *p* = 0.011), while AMY and MDt showed significant modulation in both the Cue and Outcome periods (AMY: Cue, *F*_2, 4818_ = 23.21, *p* < 0.001; Outcome, *F*_2, 4818_ = 8.34, *p* < 0.001. MDt: Cue, *F*_2, 3102_ = 4.68, *p* = 0.009; Outcome, *F*_2, 3102_ = 8.68, *p* < 0.001). The tendency was that activity in OFC increased in the Cue period, while VS showed decreasing activity during the Outcome period, and AMY’s activity weakened in both the Cue and Outcome periods. Overall, as learning progressed, OFC’s cue-related activity increased, and subcortical regions’ activity decreased.

**Fig. 2.**
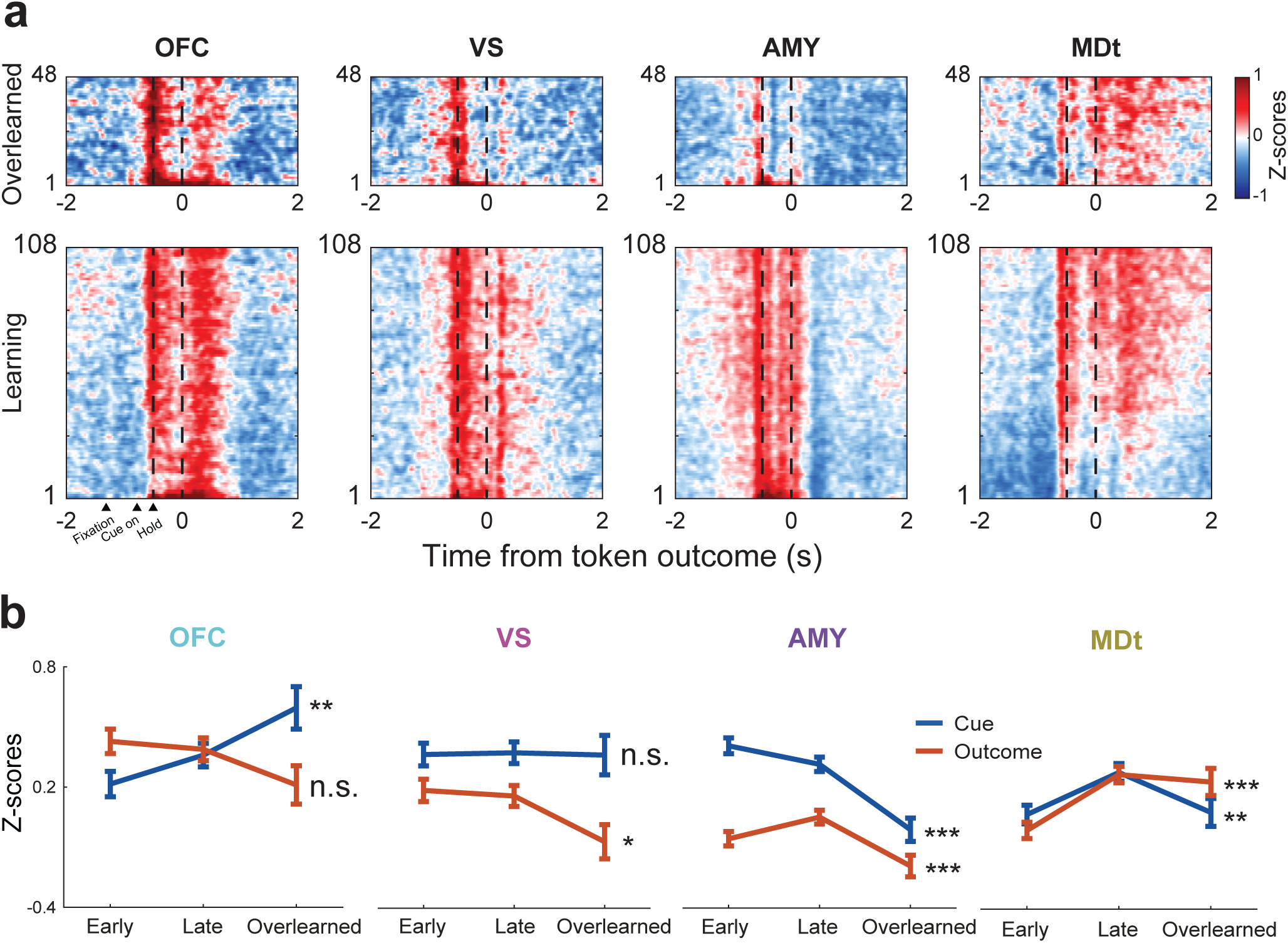
Neural activity changes with the learning process. **a,** Heatmaps of mean activity of all the neurons recorded from the OFC (n = 606), VS (n = 829), AMY (n = 1607), and MDt (n = 1035). Firing rates were Z-scored based on activity across all the trials in the same session. Warmer colors indicate higher firing rates. Activities were aligned to the onset of token outcome and ordered by the trial number in the novel (bottom) and familiar (top) blocks. **b,** Average neural responses to the Cue and Outcome periods in the Early, Late and Overlearned stages. ANOVA: n.s., not significant, \**p* < 0.05, \*\**p* < 0.01, \*\*\**p* < 0.001.

At the single-unit level, we next asked how the proportions of neurons encoding the dominant signals shifted during learning in the Cue and Outcome periods: the chosen image (Stimulus) and the change in token numbers (Δtoken). We fit single-neuron firing rates using a sliding-window multi-way ANOVA. There was a marked change in the proportion of neurons encoding the stimulus (Extended Data Fig. 1a). The response latency in OFC decreased significantly in familiar blocks, indicating that OFC neurons responded to stimuli faster, whereas the proportion in AMY decreased dramatically. Proportion of neurons encoding Δtoken remained stable during learning, with OFC having the highest proportion (Extended Data Fig. 1b). In VS and MDt, the proportion of neurons encoding stimulus and Δtoken remained stable across learning stages.

### Decoding of choice option conditions

Given that single-unit analysis cannot capture population-level features, we examined how value-related information was represented in the neural population. We first focused on decoding the value of image pairs (i.e., the stimulus condition). Because the stimuli changed every block and some were rarely chosen, we did not have sufficient trials to decode all chosen stimuli. Therefore, we conducted decoding analyses to assess the strength and latency of representations for the six choice conditions. With advancing learning, the peak decoding accuracy in all regions increased, indicating increasing discrimination of the condition (Fig. 3a). Decoding latency, defined as the earliest time points at which decoding accuracy was significantly higher than the Fixation period, shortened in all regions, showing faster recognition of different conditions (Fig. 3b).

**Fig. 3.**
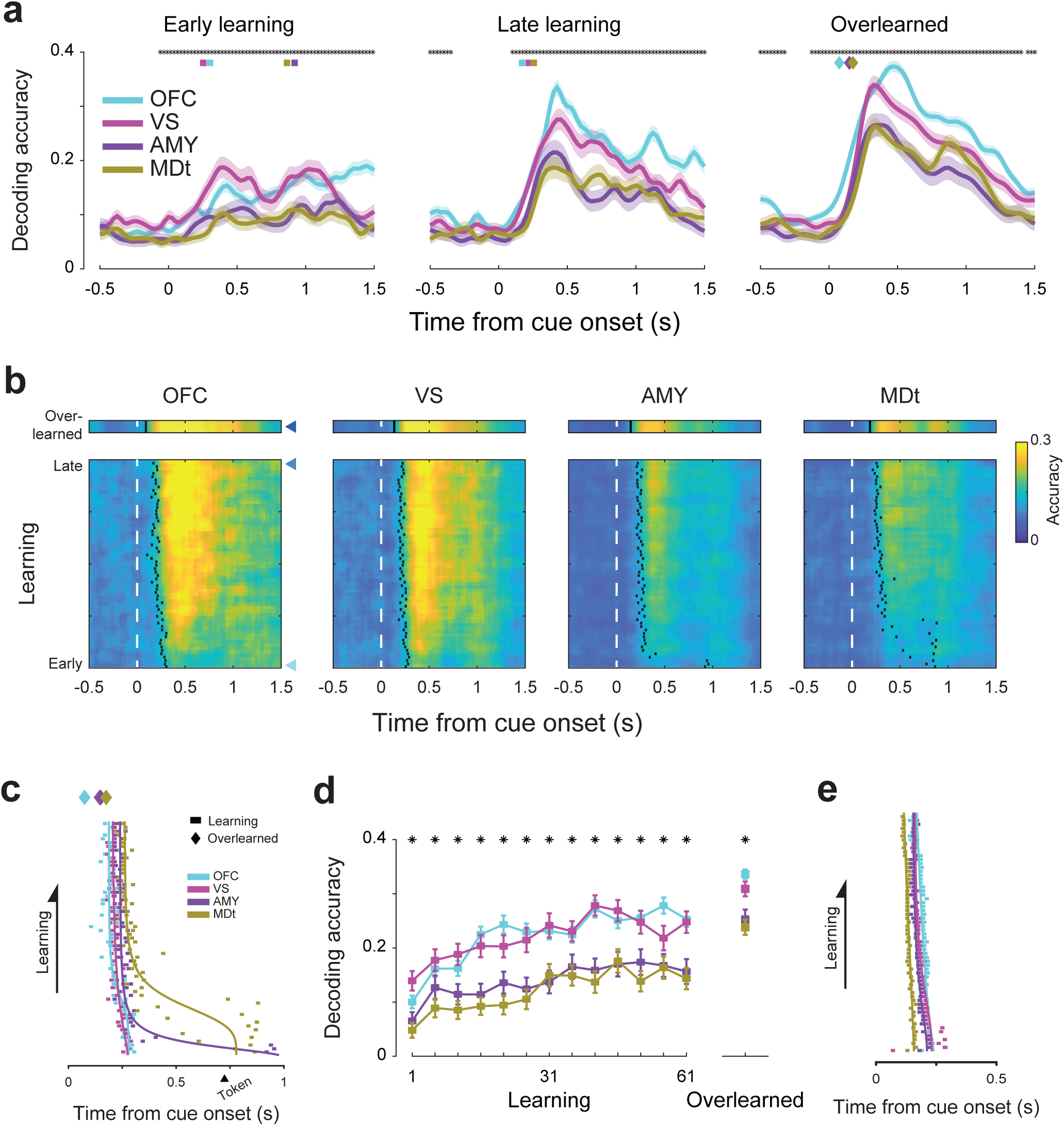
Decoding of choice option conditions. **a,** Decoding accuracy between conditions. Trials were grouped into Early, Late, and Overlearned stages. The analysis was repeated 1000 times, with 500 neurons randomly selected from each population. Shaded zones show the mean ± SD across repetitions. The black asterisks indicate a significant difference among the 4 regions (bootstrap test, *p* < 0.01). Inset squares and diamonds indicate the decoding latency for different regions. **b,** Sliding decoding using bins with a fixed number of trials (n = 48, equal to the number of trials in novel blocks). The bottom subplots correspond to novel blocks, stepped by 1 trial. Top bars correspond to familiar blocks. The black dots indicate the decoding latency. The 3 blue triangles mark Early, Late, and Overlearned stages, as shown in subplot **a**. **c,** Comparison of decoding latency among regions for conditions. Lines show the best-fit sigmoids. Diamonds on the top indicate the Overlearned stage. **d,** Mean decoding accuracy. Error bars show the mean ± SD across repetitions. The black asterisks indicate a significant difference among the 4 regions (bootstrap test, *p* < 0.01). **e,** Decoding latency of chosen direction. Lines show the best-fit sigmoids.

Learning reshaped relative decoding latency and accuracy among the 4 regions. In the early stage of learning, significant decoding first emerged in VS, whereas OFC preceded VS in the late and overlearned stages (Fig. 3c, OFC-VS, first 20 decoding chunks in novel blocks, *t*_19_ = 2.187, *p* = 0.042; last 20 decoding chunks, *t*_19_ = −2.367, *p* = 0.029). We also fit a sigmoid curve to visualize this shift (Fig. 3c). Decoding accuracy revealed a similar pattern. OFC’s accuracy was lower than VS in the early stage, but surpassed VS in the late and overlearned stages (Fig. 3d, OFC-VS, first 20 decoding chunks, *t*_19_ = −2.555, *p* = 0.019; last 20 decoding chunks, *t*_19_ = 3.595, *p* = 0.002). Overall, the decoding results showed that the strongest and earliest decoding of value shifted from VS to OFC. We also checked the decoding latency of the chosen direction as a control, where MDt showed stable, early significant decoding (Fig. 3e).

### Representation of chosen value

We next examined chosen value, the assigned value of the chosen image. To extract it from neural population activity, we performed targeted dimensionality reduction ^33^. This analysis first defined axes of task-related variables in the space of neural populations using regression coefficients from a linear model, and then projected neural activity onto the axis corresponding to the variable of interest (Fig. 4a-c). Using this approach, we isolated chosen value, which was embedded in the mixed neural representation.

**Fig. 4.**
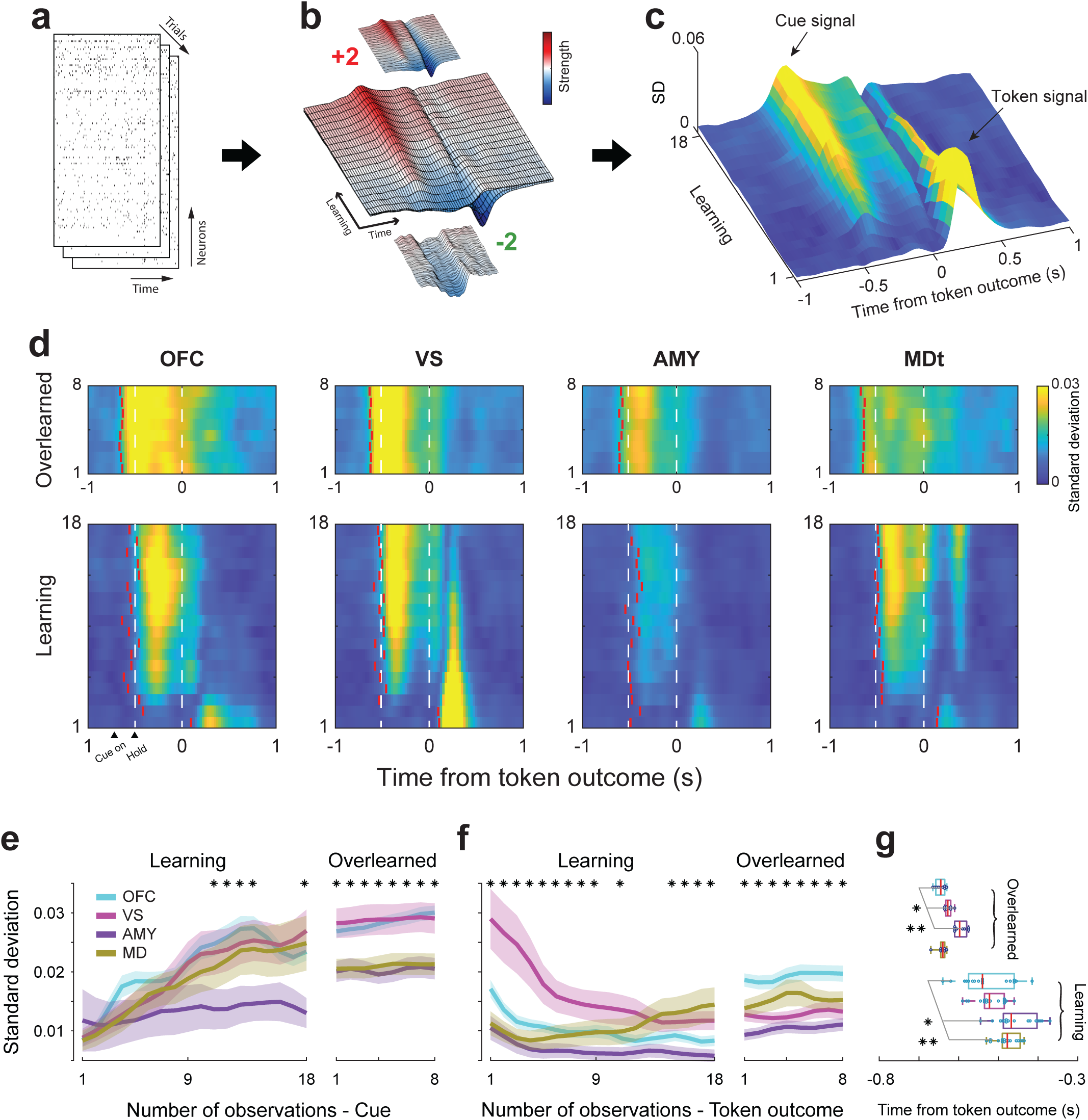
Representation of chosen value. **a-c,** Targeted dimensionality reduction and definition of encoding strength. **a**, Population activity across trials. **b,** Neural population’s representation of chosen value extracted by targeted dimensionality reduction. The X- and Y-axes represent time during the trial and learning process. The Z-axis represents the chosen value, red for positive and blue for negative. Trials are grouped by the assigned value of the chosen image. The 4 surfaces corresponding to each group are shown together in the central plot. **c**, Encoding strength of chosen value in VS. The strength is defined as the discrimination of the population representation of assigned values, quantified as the standard deviation along the Z-axis in subplot **b**. The strongest encoding of the chosen value in VS shifts from the token signal to the cue signal. **d,** Encoding strength of all regions as a function of time in the trial and learning process. Blue stands for weak encoding and yellow stands for strong encoding. Plots at the bottom and top are from novel and familiar blocks, respectively. Red bars indicate encoding latency, defined as the earliest time at which encoding strength differs significantly from baseline. **e-f,** Changes of encoding strength along the learning process. **e** for Cue period and **f** for Outcome period. Shaded areas show 95% CI. **g,** Boxplot of decoding latency. The upper 4 boxes are from familiar blocks, and the lower 4 are from novel blocks. The color of areas is the same as **e** and **f**. Wilcoxon signed-rank test: \**p* < 0.05, \*\**p* < 0.01, \*\*\**p* < 0.001, only comparisons with OFC are shown.

We wanted to ask how strongly each region encoded chosen value. The stronger a region encodes chosen value, the more distinct its responses should be for different values. We classified the trials into 4 groups of chosen value +2, +1, −1, and −2, calculated their respective mean activity in each group (Fig. 4b), and defined the standard deviation across groups (which captures the spread of the average for each value) as the encoding strength (Fig. 4c). We found that the strongest encoding appeared in the Cue period (Fig. 4e) and the Outcome period (Fig. 4f).

We found an apparent transfer of encoding time in VS, from the Outcome period to the Cue period (Fig. 4c). Early in learning, VS represented changes in token outcomes. As learning progressed, the value signal shifted gradually from tokens to images as the monkeys became more familiar with images’ value later in the block. This represents the classic transfer of TD reward prediction errors from outcome to cue ^34,35^, further supported by the overlearned data. In familiar blocks, the monkeys remembered the image values, so learning was no longer involved. As a result, value information was carried by images rather than tokens (Fig. 4d top row). The pattern was absent or not as strong in the other 3 regions (Fig. 4d, bottom row). In OFC, during the Outcome period, encoding strength rapidly declined from a moderate level during early learning. During the Cue period, encoding strength increased during novel blocks and was maintained and extended in time in familiar blocks. For AMY, encoding in the Cue period was stable but low in novel blocks and significantly rose in familiar blocks. MDt showed increasing encoding strength in novel blocks during both the Cue and Outcome periods. But in familiar blocks, the two periods diverged: the Outcome period showed greater strength, while encoding in the Cue period weakened.

We next compared their encoding latencies (Fig. 4g). No significant difference between OFC and VS was observed in novel blocks, but OFC encoding was earlier than VS in familiar blocks (Wilcoxon signed-rank test, *p* = 0.031). In short, early in learning, the VS drove value updates, but as learning progressed, the OFC became the primary encoder of the chosen value.

### Unique encoding of state value in OFC

Considering its significance above, we further investigated how the OFC encodes state value. State value is the expected future reward a monkey can obtain starting from a given state ^36,37^. It reflects the subjective value of that state to the animal. For example, the Fixation period in the late learning stage has a higher value than in the early learning stage, because monkeys are more likely to receive rewards due to experience with the cues. To estimate state value, we constructed a POMDP model ^38^ based on the R-W model’s value in each trial (Fig. 5a-b), but also including additional factors related to the number of accumulated tokens and time since the last cashout. It divided a trial into several epochs (ITI, Fixation, Cue, Outcome, Cashout) and computed their values (Fig. 5b). Specifically, our model accounted for learning effects on state value.

**Fig. 5.**
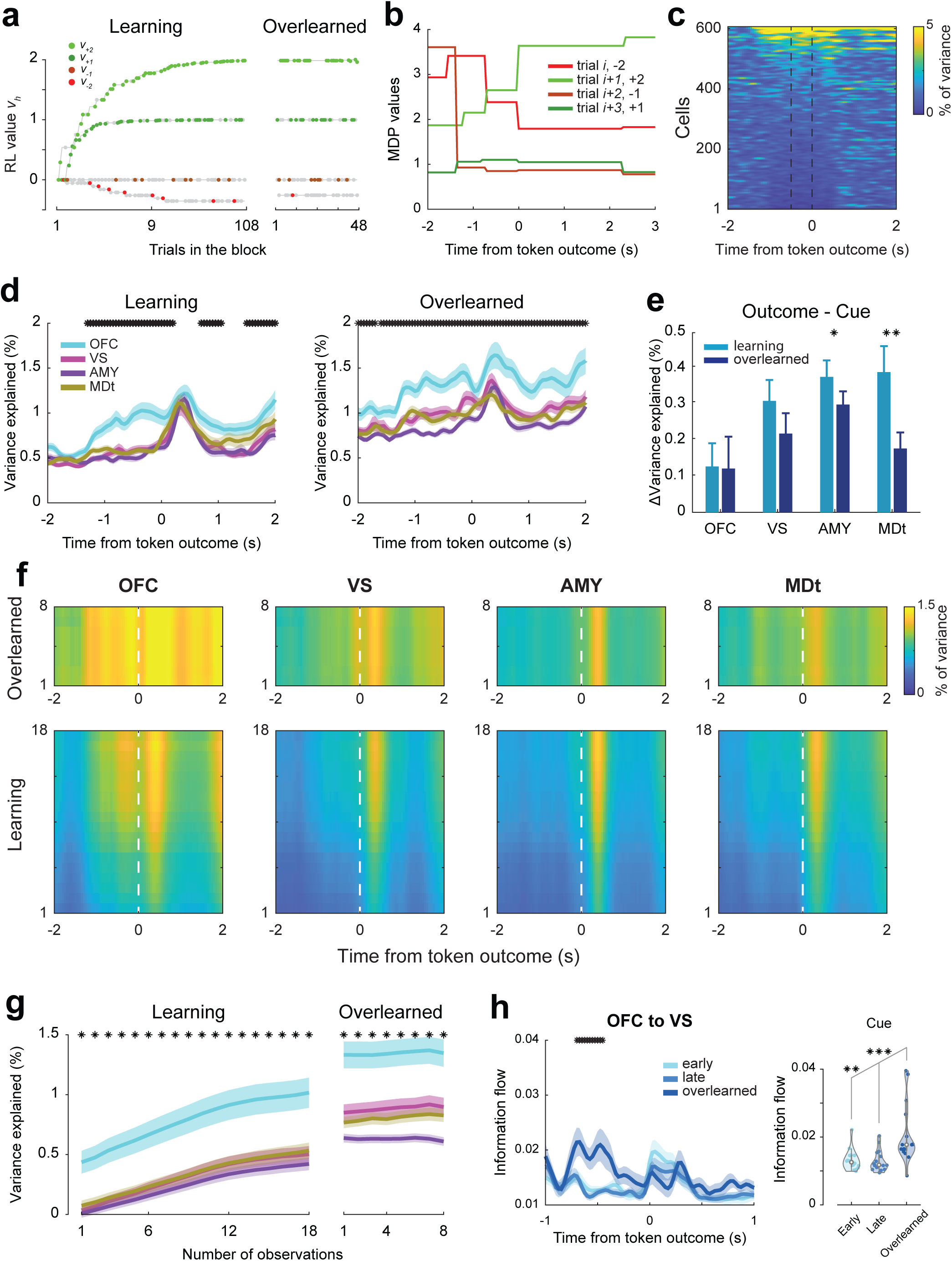
Unique encoding of state value in OFC. **a,** The updating of images’ value in an example block from the RW model. Each dot represents one image presented on the screen in each trial; color dots represent chosen images, and gray ones were not chosen. The resolution of the RW value was at the trial level. **b,** State values in 4 continuous example trials. Trials were split into 5 sections, each assigned a value estimated by the POMDP model. The resolution of state value was at the sub-trial level. **c,** Variance of neural activity explained by state value in OFC, familiar blocks. Neurons were sorted by the mean variance explained in the trial. **d,** Proportion of variance explained by state value in the 4 recorded regions along the axis of time in the trial. Learning stands for novel blocks, and Overlearned stands for familiar blocks. Shaded areas indicate SEM, and black asterisks indicate significant differences among the 4 regions (ANOVA, *p* < 0.01). **e,** Difference of variance explained between the Outcome and Cue periods, *t*-test: \**p* < 0.05, \*\**p* < 0.01, \*\*\**p* < 0.001. **f.** Heat plot of variance explained along the axis of time in trial (X-axis) and learning process (Y-axis). The bottom subplots were from novel blocks, and the top subplots were from familiar blocks. The numbers on the Y-axis indicated how many times monkeys had seen a specific image pair. Brighter regions indicate more explanatory power of state value to neural activity. **g.** Variance explained of the 4 regions along the learning axis. Each value is the mean over the period from 1000 ms before the token outcome to 500 ms after. Black asterisks indicate a significant difference among the 4 regions (ANOVA, *p* < 0.01). **h,** Information flow from OFC to VS in the 3 learning stages. Colors from light to dark correspond to early, late, and learned, respectively. Black asterisks indicate a significant difference among the 3 stages (ANOVA, *p* < 0.01). Violin plots show the distribution of information flow in the Cue period (700-400 ms before token outcome). Wilcoxon signed-rank test: \**p* < 0.05, \*\**p* < 0.01, \*\*\**p* < 0.001.

We fit the firing rates of neurons with a multivariate linear model that included state value as a predictor, and used the percentage of variance explained by state value as a measure of encoding strength. We first examined the distribution of encoding strength within the neural population (Fig. 5c). Only a small proportion of neurons encoded state value strongly. We further investigated the encoding strength of each region (Fig. 5d-g). Fig. 5d shows the mean encoding of state value across a trial in the novel (Learning) and familiar (Overlearned) blocks. We expect that as learning progresses, the state value propagates backward from the Outcome to the Cue, Fixation, and the inter-trial interval (ITI), as monkeys become more certain about the outcomes they will receive and the choices they will make.

As expected, state value encoding increased in all regions after learning, in the Outcome and also Cue periods (Extended Data Fig. 2f). For AMY and MDt, the encoding difference between the Outcome and Cue periods decreased after overlearning (Fig. 5e), suggesting an extra gain of state value in earlier periods compared to token outcome. However, OFC uniquely encoded state value with greater strength and earlier timing (Fig. 5d-g). In novel blocks, encoding in OFC was significantly stronger than in other regions from the Fixation period to the Outcome period (Fig. 5f). In familiar blocks, this advantage expanded to even earlier stages in the ITI. We also examined the encoding of the difference in state value across trial periods, and OFC was the strongest encoder as well (Extended Data Fig. 2, a-e). Specifically, OFC exceeded the other 3 regions in encoding the difference between Fixation and ITI. These results suggest that OFC uniquely encodes early state value, a key distinction from chosen value and token outcomes.

Having observed that state value first appeared in OFC, we next asked how it propagated across the 4 regions in the trial and how the information flow changed with learning (Extended Data Fig. 3). We performed Granger causality analyses to quantify information flow for state value (see Method). Flow from OFC to VS in the Cue period remained low in novel blocks, but rose significantly in familiar blocks (Fig. 5h, Wilcoxon signed-rank test, Early - Late, *p* = 0.278; Early - Overlearned, *p* = 0.015; Late - Overlearned, *p* = 0.004), accompanied by strengthened state value encoding in OFC (Fig. 5f). In contrast, flow from VS to OFC didn’t change significantly, indicating a unidirectional enhancement (Extended Data Fig. 3a). This indicates a shift in the leading value from the cue value in VS to the state value in OFC.

## Discussion

We examined the single-unit, population, and network representation of value information across the ventral cortico-striatal-thalamo-cortical network. Our findings reveal a dynamic functional reorganization of this circuit as learning progresses from early to late to overlearned. Specifically, we observed that while the VS was crucial for rapid updating of stimulus value during early learning, the OFC showed stronger and earlier value representation, and increased state value representation during overlearning. This results in a significant shift of the information processing center from subcortical to cortical structures, providing a direct challenge to traditional “cortical-to-subcortical” shift hypotheses ^17,39^.

### Distinguishing learning from overlearning

Previous studies have examined learning across different time scales by collecting data over a long-term training history ^28,40^. This addresses the learning process at a single-day level. However, it compares the coding of learning across different groups of neurons. To address the neural basis of different learning stages within a single session, we employed both novel and familiar blocks in the task, allowing us to probe the same neurons across distinct learning stages.

Previous studies suggest a shift from goal-directed to automated behavior after sufficient learning ^5,6,16^. Our behavioral results were consistent with this perspective, where monkeys’ choice preferences evolved rapidly in novel blocks (goal-directed learning) but remained stable in familiar blocks (automated behavior without learning). Different learning stages vary in their reliance on neural plasticity and the speed of signal updates. While prior work has separately implicated the OFC, VS, and AMY in value processing ^15,33^, their specific roles within a broader network remain unclear, especially during the transition from early acquisition to overlearning ^41^.

### The ventral striatum encodes early value updates

Although value updating has been attributed to the VS, previous evidence was controversial. fMRI studies in humans reported RPE-related BOLD signal in VS ^12,42–44^, but single-unit recording suggests VS acts like a “critic” but not simply a RPE-encoder ^10,13^. This may be due to the fact that VS is not simply encoding RPE, but an integration of multiple values ^45^ and temporal information ^46^. To isolate the RPE-related signal from this mixed information, we applied targeted dimensionality reduction to the neural populations. We found that the VS is primarily involved in updating stimulus values during early learning. In early learning, the VS showed earlier and stronger encoding of stimuli and outcomes than the other regions. The major change in VS was that the encoding of values shifted from the reinforcement feedback to its corresponding stimulus. This is consistent with the classic view of the VS as a primary recipient of dopaminergic temporal-difference reward prediction errors (RPE) ^35^ and its role in rapid associative learning ^47–50^.

We also found that the expected value representation in VS is limited to the stimulus period. This implies that information abstraction in VS is related to sensory events, rather than explicitly holding information online. In early learning, as learned information is simple, rapid, and model-free, VS is dominant in the network. But once the value is assigned and the association is consolidated, the computation shifts to OFC.

### OFC contributes to the encoding of state value

As the monkeys transitioned from early to late learning and then to the overlearned stage, we observed that both stimulus and value processing shifted from VS to OFC, characterized by higher decoding accuracy and shorter information latency. This finding contrasts with the suggestion that the cortex should become less involved late in learning ^31^. Previous findings suggest the role of OFC is to encode state information ^30,36^, and we found OFC was the main encoder of state value among the recorded regions. In late learning in novel blocks, we observed increased information in OFC during stimulus encoding, not during value encoding. In this period, OFC has learned the state determined by the presented images, but these states are not firmly associated with value. Only after extensive training does the OFC show the strongest and earliest representation of value, as indicated by condition decoding and the encoding latency of the chosen value. Our results on information flow also support this, showing that flow from OFC to VS is only enhanced in familiar blocks, not in late novel blocks. We suppose the meaning of this flow is to use state value in OFC to guide later action selection in the basal ganglia, and this requires state value in OFC to be reliable ^14^, which is present during overlearning.

In conclusion, the shift of information processing from VS to OFC corresponds to the advancement from stimulus value to state value ^51^, which aligns closely with computational frameworks of frontostriatal function. While the VS handles localized, model-free value predictions based on explicit cues ^13,52^, the OFC constructs a latent cognitive map of the environment ^36^. Consequently, the shift from VS to OFC represents an elevation from simple, flat associative tracking to a structured, state-based representation of the task space necessary for model-based reinforcement learning ^3,53^.

### The role of the ventral cortico-striatal-thalamo-cortical network in learning

Our findings reveal that the ventral cortico-striatal-thalamo-cortical network operates as a dynamic system that redistributes processing as expertise is gained. During early learning, the network operates in a “bottom-up” manner, driven by subcortical dominance of the VS and AMY to rapidly encode reinforcement signals and allocate reinforcement ^4,15^. Within this stage, value updating in the VS is limited to the immediate outcome. Here, the VS handles model-free value predictions ^13,52^, shifting the encoding of value from feedback to the corresponding stimulus ^35^. These rapid subcortical signals serve as guiding inputs, relayed via the MDt to slower-learning cortical systems ^54^. Once stimulus value is consolidated, the computational burden shifts. This transition begins in late learning, where a cortical advantage emerges in the OFC for stimulus encoding. The OFC has mapped the task states, but they are not yet firmly associated with value ^30^.

Ultimately, after overlearning, the network hierarchy reverses, shifting the information processing center from subcortical to cortical structures ^55^. The center of both stimulus and value processing shifts from the VS to the OFC. During overlearning, the OFC acquires state value encoding and emerges as the main node ^56^. Consequently, communication becomes “top-down”: enhanced information flow runs exclusively from the OFC to the VS in familiar blocks to guide subsequent action selection in the basal ganglia ^14^. This shift from VS to OFC corresponds to an advancement from stimulus value to state value, lifting the abstraction level from simple model-free associative learning to structured, model-based reinforcement learning ^3,36,57^.

## Methods

### Subjects

The experiments were performed on two adult rhesus macaques (*Macaca mulatta*), a 10 kg male and a 7.5 kg female, between 8 and 10 years old. The monkeys were housed in pairs whenever possible and fed on a 24-hour schedule. On testing days, water intake was controlled, requiring the subjects to earn juice rewards through task performance. Conversely, on non-testing days, water was provided *ad libitum*.

### Ethics

Experimental procedures for all monkeys followed *the Guide for the Care and Use of Laboratory Animals* and were approved by the National Institute of Mental Health Animal Care and Use Committee.

### Experimental Setup

Monkeys were trained to perform a saccade-based two-armed bandit task. Stimuli were presented on a 19-inch LCD monitor situated 40 cm from the monkeys’ eyes. During training and testing, the monkeys sat in a primate chair with their heads restrained. Stimulus presentation and behavioral monitoring were controlled by MonkeyLogic ^58^. The eye movements were monitored at 400 fps using a Viewpoint eye tracker (Arrington Research, Scottsdale, AZ) and sampled at 1 kHz. A fixed amount of apple juice was delivered via a pressurized plastic tube, gated by a solenoid valve, on rewarded trials.

### Task Design

The task was based on our previous study ^32^. Each block used a set of 4 images corresponding to the values +2, +1, −1, and −2. The monkeys obtained more tokens by choosing images associated with larger values. Choosing one of the images led to gaining or losing a corresponding number of tokens with a probability of 75%, and no token change for the other 25% (Fig. 1b). Also, token numbers could not be negative, so choosing a loss image when there were no accumulated tokens had no effect.

To complete a trial successfully, the monkey first acquired and held central fixation for 400–600 ms. Then, the computer randomly selected 2 of the 4 images and displayed them on the screen. The animal made its selection by saccading to one of them. The unchosen image disappeared when the monkey reached the target. Saccade fixation was maintained on the chosen image for 500 ms. After that, the chosen image disappeared, and the token number was updated accordingly. Tokens accumulated across trials and were cashed out for juice every 4 to 6 trials, with the interval randomly selected. At cash-out, the animals received 1 drop of juice per token. When each drop of juice was delivered, one token was removed from the screen. Up to 12 tokens were accumulated and displayed on the screen across trials.

Task condition is defined as the values of image pairs, therefor there are 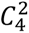 = 6 conditions. Blocks were classified into 2 types: novel and familiar. Each session comprises 9 novel blocks and 3 familiar blocks. In the novel blocks, we introduced 4 images at the beginning, and the monkey had no experience with their value, which had to be learned by choosing and observing the outcomes. A novel block had 108 trials (6 conditions × 2 counterbalanced for left and right sides × 9 repetitions). Familiar blocks relied on the training period before the experiment. During training, we repeatedly presented image sets to the monkeys, thereby providing prior knowledge of the images’ values before the experiment. In familiar blocks, these image sets were presented again, allowing monkeys to rely on their experience rather than learn from scratch. A familiar block had 48 trials (6 conditions × 2 counterbalanced for left and right sides × 4 repetitions). The conditions within each block were presented in a pseudo-random order. The animals saw each condition twice, once on the left and once on the right, every 12 trials, before seeing it a third time.

### Surgical procedures

To prepare for the study, a titanium headpost and a 25 × 35 mm recording chamber were surgically implanted into each monkey, enabling vertical grid access to the OFC, VS, AMY, and MDt ^33^. 3.0 T T1-and T2-weighted MRI scans were used to plan and confirm the precise placement of the chambers. Small burr holes were drilled above the specific target regions. Within the installed grid, guide tubes were inserted through these holes, lowered to a position 1–4 mm above the target regions, and secured to the grid with adhesive. Every 3–5 recording days, the guide tubes were replaced and shifted to adjacent locations, with their new positions verified via subsequent MRI scans. All surgical procedures were conducted under sterile conditions while the subjects were under anesthesia.

### Neurophysiological Recordings

Once the monkeys recovered from surgery, neurophysiological recordings commenced. On each recording day, linear electrode arrays (V-probe, Plexon Inc, Dallas, TX) were lowered to target regions through the guide tubes. Specifically, 32-channel probes with 150 μm spacing were utilized for the VS and MDt, while 64-channel probes with the same spacing were used for the OFC and AMY. A 4-channel micromanipulator (NAN Instruments, Nazareth, Israel) attached to the chamber was used to advance the probes to their targets. Neurophysiological data were recorded using a 512-channel Ripple Grapevine System (Ripple, Salt Lake City, UT), with the spike threshold set to 4.0 times the root-mean-square (RMS) of the baseline signal. To synchronize the data, behavioral markers from MonkeyLogic and eye-tracking signals from Viewpoint were integrated into the Ripple system. The extracellular signals were high-pass filtered (1 kHz cutoff) and digitized at 30 kHz to capture single-cell activity, with spikes subsequently sorted offline using Wave_clus 3 ^59^.

### Choice behavior

Each block had six conditions, pseudo-randomly arranged in the blocks. The number of observations in each condition is regarded as progress of learning, which increases from 1 to 18 in novel blocks, and from 1 to 8 in familiar blocks. We first computed the fraction of choosing the image associated with the higher value (correct rate, CR) in each condition to evaluate choice behavior. We fitted monkeys’ behavior with the RW model (Fig. 1d).

To compare performance across stages, particularly late novel and familiar blocks, we conducted a two-sample proportion test. Trials were divided into 3 groups: Early was the first 48 trials in novel blocks, Late was the last 48 trials in novel blocks, and Overlearned was all trials from familiar blocks. The number 48 was set to balance the number of trials per group, which was also used in the neural analyses. We conducted a pairwise comparison among the 3 trial groups using the chi-squared test for proportions.

### Neural activity and proportion of responsive neurons

To analyze neural activity changes resulting from learning, we extracted two time periods from each trial: Cue and Outcome, during which firing rates were highest. The Cue period is a 500-ms window at 700-200 ms before the token outcome, beginning approximately at cue onset. The Outcome period was set to 0-500 ms after the token outcome. The mean firing rate for each neuron was computed in each time window and each learning stage, and ANOVAs were performed to check the main effect of learning stage on firing rate in each recorded region.

To identify neurons that respond to different task components at different times during trials, we fit a sliding-window multi-way ANOVA model to spike counts computed in 200 ms bins (Extended Data Fig. 1). The bins were aligned to the time of the token outcome and advanced by 50 ms. Factors in ANOVA included the number of tokens on the screen (#token, may change after the choices), the change of token numbers (Δtoken), the drops of juice delivered at juice Outcome period (#juice), image pairs with different value combinations presented (condition, e.g., +2 vs. −1), the order of blocks (blockID), the assigned value (cValue), and direction of the chosen image (cDir, left or right), and the identity of the chosen image (Stimulus). The factor blockID was used to remove non-stationarity due to drift. Significant encoding of each factor at each time bin was evaluated at *p* < 0.05. A neuron that showed a significant response to a factor in no less than three contiguous bins in the statistics was considered to be responsive to this factor. We then examined the proportion of responsive neurons across different brain regions, using the binomial test to determine whether each region had a proportion significantly above chance (5%).

### Decoding analysis

We decoded the trial condition (values of the image pair) from the firing rates of neurons in different regions. We balanced the number of trials used in different learning stages. For each session, 3 familiar blocks and 3 novel blocks (randomly chosen from 9) were used. For each decoding round, we used continuous 48 trials from 3 blocks of the same type (e.g., the 1st through 48th trials in 3 novel blocks for Early learning). We generated 1,000 bootstrap samples, each consisting of 500 neurons randomly drawn from the dataset. Then, decoding was obtained with leave-one-out cross-validation.

Decoding latency was defined as the earliest time point at which decoding accuracy was significantly higher than during the Fixation period. The Fixation period was 500-0 ms before cue onset, and the accuracy baseline in a bootstrap sample was the mean decoding accuracy for this period. The criterion of significant was that decoding accuracy was higher than the baseline in over 99% bootstrap samples. We also conducted the same analysis when decoding the chosen direction.

Because of the 48-trial window, we obtained 61 windows from novel blocks and 1 from familiar blocks. Decoding latency of 61 windows in novel blocks was fit by a sigmoid curve. We also checked changes in decoding accuracy across these time windows, using the mean accuracy during 200-500 ms after cue onset to represent the cue information before the token outcome.

### Targeted dimensionality reduction

This method was applied to extract the represented value of the chosen image from the neural population activity. It is based on a series of multivariate linear models. The dependent variable, neural activity, was the z-scored firing rate, computed per neuron at the session level. We employed 8 task-relevant independent variables, as described in the section on the Proportion of responsive neurons. The linear model can be formalized as:

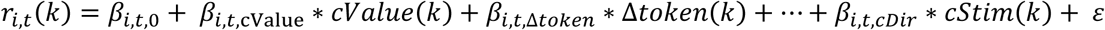

where *r_i,t_* (*k*) is the z-scored response of neuron *S* at time *t* and on trial *k, cValue*(*k*) is the assigned value of the chosen image on trial *k*. The regression coefficients, *β_i_*_,*t*,*f*_, describe how much the trial-by-trial firing rate of neuron *S*, at a given time *t* during the trial, depends on the corresponding task variable *f*. *β_i_*_,*t*,0_ is the intercept. For a given time bin *t*, we regard the population activity as a vector, each component of which corresponds to a single neuron. Then the linear model can be written in vector form:

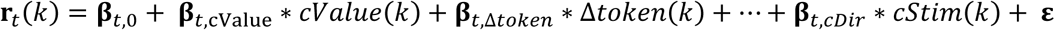

**β***_t_* can be considered as axes of task-relevant variables. By projecting **r***_t_* to the direction of **β***_t_*_,cValue_, we obtain a represented chosen value in the neural population ^33^. Here we do this by dot product:

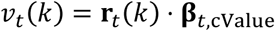

where *v_t_*(*k*) is the represented chosen value at time bin *t* in trial *k*.

To map the represented chosen value to encoding strength, we categorized all trials by the actual chosen value, yielding 4 groups: +2, +1, −1, and −2. For each group, each number of observations of conditions and each time bin, we computed the mean of represented chosen value (Fig. 4b). Then, we calculated the standard deviation on the group dimension using 4 data points, which was defined as encoding strength of chosen value, as a function of number of observation (learning process) and time in the trial (Fig. 4c, d).

All operations above were done using the bootstrap method. For each of 1000 iterations, we randomly sampled 500 neurons. Fig. 4b-d displayed the mean of 1000 samples. Fig. 4e-f showed the mean and 95% CI. Here, the Cue period was 500–0 ms before the token outcome, and the Outcome period was 100–600 ms after the token outcome.

We also checked the encoding latency of chosen values. We set the criterion for significant encoding as the mean plus 5 standard deviations of baseline, and encoding latency was the earliest time at which the mean encoding strength across bootstrap samples exceeded the criterion. Red bars in Fig. 4d indicate encoding latency, and Fig. 4g shows their distribution along the time course of a trial.

### Rescorla-Wagner model

We used a variant of the RW model, as was previously used to model the tokens task ^32^.

We fit a RW value update equation given by the following:

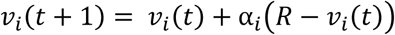

where the variable *v_i_* is the value estimate for cue option *i* that was chosen on trial t, *R* is the change in the number of tokens that followed the choice in trial *t*, and α*_i_* is the cue-dependent learning rate parameter.

The value computed above was then used to compute choice probabilities for each cue pair using the softmax function:

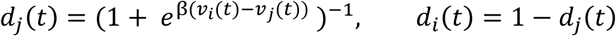

where β, is the consistency choice parameter, fit across all six cue conditions, and *i* and *j* are the two choice options. We then maximized the likelihood of the animal’s choices, *D*, given the parameters:

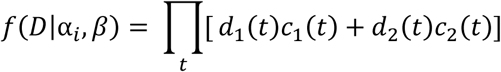

where *d*_1_(*t*) is the choice probability value for option 1 on trial *t*, and *c*_1_(*t*) and *c*_2_(*t*) are indicator variables that take on a value of 1 if the corresponding option was chosen and 0 otherwise. This model was fit across blocks within each session for each monkey, yielding a single set of fit parameters per session.

### Partially observable Markov decision process model

State values were computed using a Markov decision process (MDP) model adapted from our previous work ^38^.

The MDP model computes the value of each task state, defined as the sum of immediate rewards and the expected discounted sum of future rewards from that state. This is referred to throughout as the SV or the utility of the state *u*(*s_t_*). The state *s_t_* is a function of five task features: the number of accumulated tokens (NTk), the number of trials since cash-out (TSCO), the task epoch (TE), and the number of observations of a cue pair within a block (NObs). All pairwise combinations of features satisfying NTk ≤ 12 were included as valid states, with a maximum of 12 total tokens consistent with the six-trial maximum duration until cashout. The state value was computed using value iteration ^60^ applied to the following Bellman optimality equation:

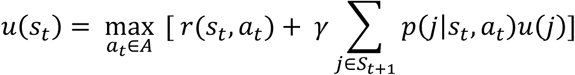

where *s_t_* is the state, *a_t_* is the action taken at that state, *u*(*s_t_*) is the SV, *r*(*s_t_*, *a_t_*) is the immediate reward, γ is the discount factor, *p*(*j*|*s_t_*, *a_t_*) is the transition probability to future state *j* and *S_t+1_* is the set of immediate future possible states from state *s_t_* if one takes action *a_t_*. The summation over future states (*j*) is the expected future utility, taken across the transition probability distribution *p*(*j*|*s_t_*, *a_t_*). As in the original published model, the discount factor was γ = 0.999. Value iteration was run until the policy and state values converged, which required approximately 100 iterations.

All state features and transitions were identical to those in ^38^. The transition probabilities from fixation to any of the six cue states were modeled as *p*(cues) = 1/6. Cash-out transition probabilities were *p*(cashout) = 0 for TSCO 1–3, *p*(cashout) = 0.33 for TSCO = 4, *p*(cashout) = 0.50 for TSCO = 5, and *p*(cashout) = 1.0 for TSCO = 6. As in the original model, transition probabilities for the token outcome given a choice (*p*(ΔNTk | choice)) were derived from the RW model fits to each monkey’s choice behavior (see RW model section). A separate MDP was fit for each monkey using its corresponding transition probabilities.

State values were extracted for all trials and epochs using the MDP fits for each monkey. This produced a table of states such that the value of each state was:

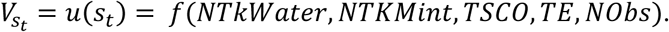

To evaluate the effect of state values on firing rate, we fit another multivariate linear model. Based on the above 8-variable model, we combined all state value-related variables and obtained a 5-variable linear model, in which independent variables are state value, direction of the chosen image (cDir), the order of blocks (blockID), identity of the chosen image (Stimulus), and whether the Δtoken is the assigned value of chosen image (isStochastic). We used the percentage of variance explained by state value as the measure of encoding strength (Fig.5).

To assess the encoding of state value changes, we modified the 5-variable model by splitting state value into differences across contiguous trial sections. A trial included at most 5 sections: ITI, Fixation, Cue, token Outcome, and juice Cashout, and differences of neighboring sections were used as factors. The percentage of variance explained by state value differences is shown in Extended Data Fig. 2.

### Information flow analysis

We used Granger causality analysis to measure the flow of information across the recorded regions based on a linear model ^61^:

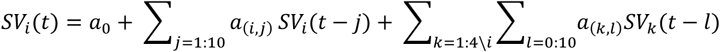

Here, *SV* represents the state values, *i* represents the output region, *j* represents the lagged bin number ahead of time *t* in the output array, *k* represents the input regions, *l* represents the bin number ahead of time *t* in the input array. SV was computed from −1 to 1 seconds relative to the token outcome, in 25 ms bins, advancing by 25 ms. Note that we are predicting the future information, *SV_i_*(*t*), using past information on the same region, 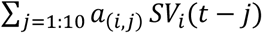 and current and past information in other regions, 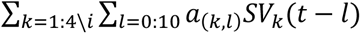. This is called the Full model, which includes all regions. We computed the variance of the prediction error of the full model, denoted as *Var_full_*.

When we tested for the effect of one region on another, we dropped the region under consideration from the sum and compared the prediction of *SV_i_*(*t*). This is called the Partial model, with a region dropped off. We then had *Var_partial_* as that of the full model. Then, the information flow of SV in this time bin was:

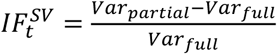

## Data availability

The datasets supporting the current study will be publicly available as of the date of publication.

## Code availability

All original codes will be publicly available as of the date of publication.

## Acknowledgments

This work was supported by the Brain Science and Brain-like Intelligence Technology -National Science and Technology Major Project under Grant 2021ZD0204300 to X. L., BBRF NARSAD Young Investigator Award (30892) to H. T., and Intramural Research Program of the National Institute of Mental Health (ZIA MH002928) to B. B. A.

## Author contributions

Conceptualization, H.T., and B.B.A.; methodology, H.T., R.B., and B.B.A.; investigation, D.X., D.B., R.B. and H.T.; visualization, D.X., D.B., B.B.A. and H.T.; writing—original draft and review & editing, D.X., X. L., B.B.A. and H.T; funding acquisition, X. L., H.T. and B.B.A.; resources, H.T., and B.B.A.; supervision, X. L., B.B.A. and H.T.

