## Supplemental Files for "Long-term Learning Induces Plastic Changes in Frontostriatal Circuits"

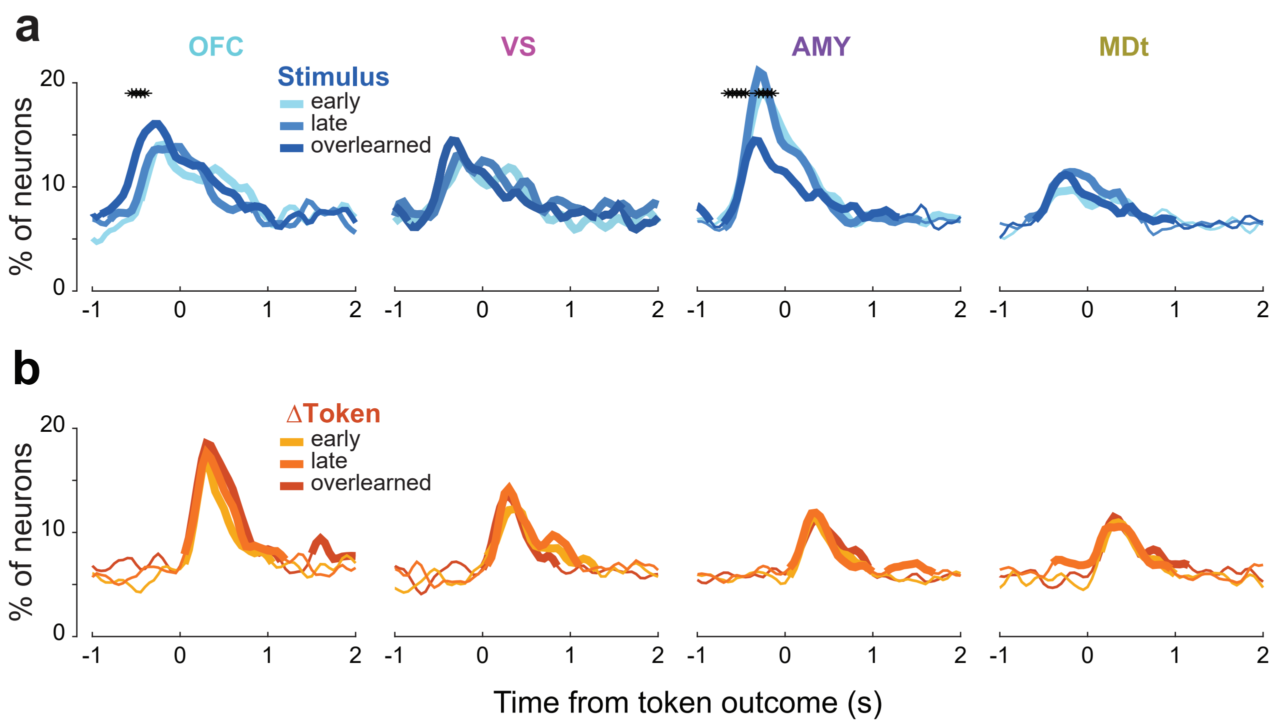


Extended Figure 1

### Extended figure legends

#### Extended Data Fig. 1 | Proportion of neurons encoding stimulus and Δ token

Proportion of neurons encoding Stimulus (**a**) and Δtoken (**b**). Thick lines indicate a significant difference from chance level (binomial test, $p < 0.01$). Black asterisks indicate a significant difference among the 3 learning stages (chi-squared test, $p < 0.05$).

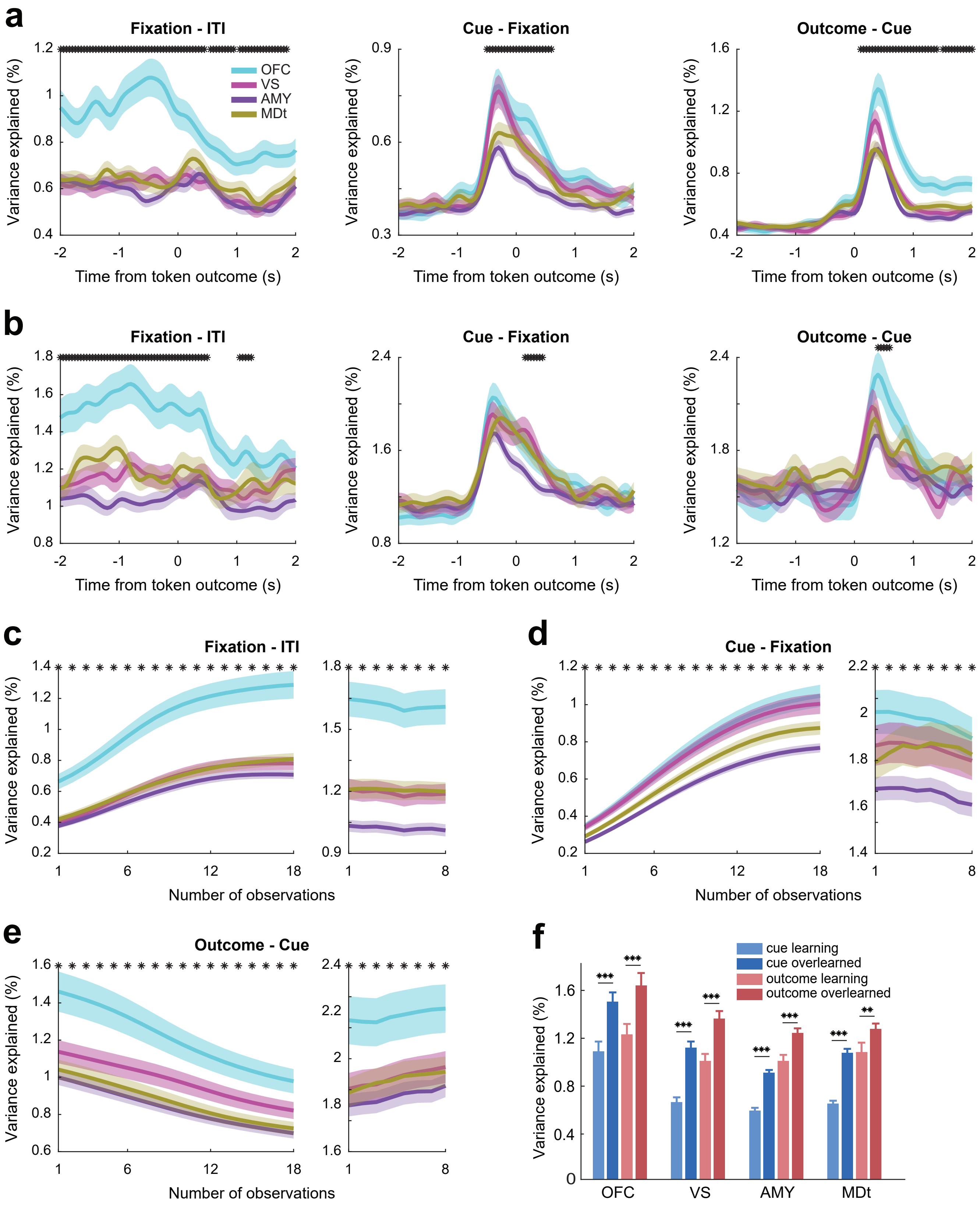


Extended Figure 2

#### Extended Data Fig. 2 | Neural activity variance explained by state value changes

**a,** Variance explained by state value changes in novel blocks. Titles indicate sections used to compute state value differences. Shaded area shows the SEM. Black asterisks indicate a significant difference among the 3 stages (ANOVA, $p < 0.01$). The same applies to other subplots.

**b,** Variance explained by state value changes in familiar blocks.

**c-e,** Variance explained by state value changes along the learning process. Left subplots show the novel blocks, and right subplots show the familiar blocks. Each value is the mean over the period from 1000 ms before the token outcome to 500 ms after. Black asterisks indicate a significant difference among the 4 regions (ANOVA, $p < 0.01$)

**f,** Variance explained by state value in Cue and Outcome periods, during learning and overlearned stages. *t*-test: *$p < 0.05$, **$p < 0.01$, ***$p < 0.001$.

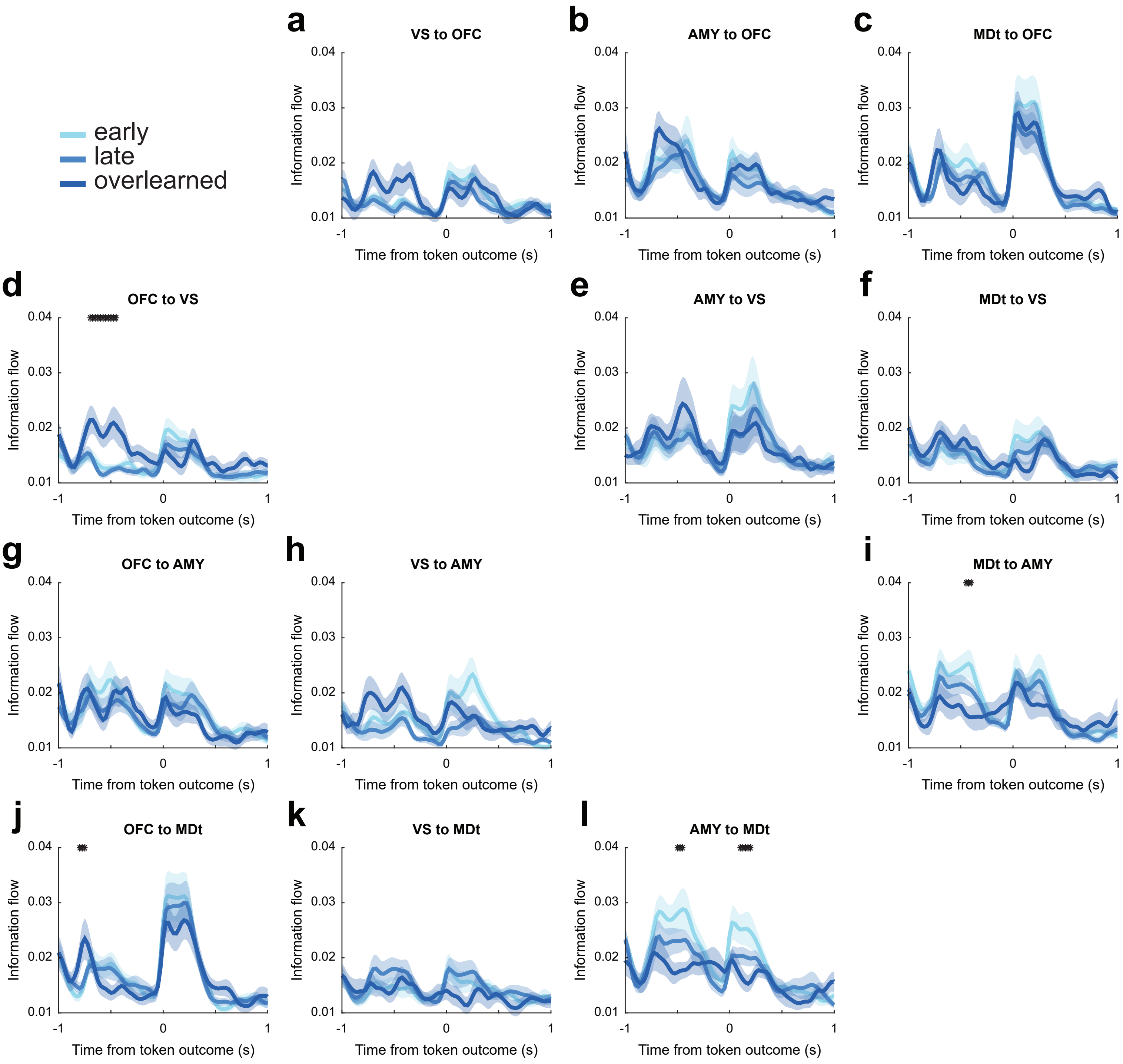


Extended Figure 3

#### Extended Data Fig. 3 | Information flow of state value among recorded regions

Each subplot shows the state value information flow along the edge specified by the title. Shaded area shows the SEM. Black asterisks indicate a significant difference among the 3 stages (ANOVA, $p < 0.01$).
